# Structural and Genetic Determinants of Convergence in the *Drosophila* tRNA Structure-Function Map

**DOI:** 10.1101/2020.07.24.220558

**Authors:** Julie Baker Phillips, David H. Ardell

**Affiliations:** Quantitative and Systems Biology Program, University of California, Merced, CA 95343; Department of Biology, Cumberland University, 1 Cumberland Square, Lebanon, TN 37087; Department of Molecular and Cell Biology, University of California, Merced, CA 95343

**Keywords:** Class-Informative Feature (CIF), ion-binding pocket, parallel substitutions, convergent evolution, structure-function map

## Abstract

The evolution of tRNA multigene families remains poorly understood, exhibiting unusual phenomena such as functional conversions of tRNA genes through anticodon shift substitutions. We improved FlyBase tRNA gene annotations from twelve *Drosophila* species, incorporating previously identified ortholog sets to compare substitution rates across tRNA bodies at single-site and base-pair resolution. All rapidly evolving sites fell within the same metal ion-binding pocket, that lies at the interface of the two major stacked helical domains. We applied our tRNA Structure-Function Mapper (tSFM) method independently to each *Drosophila* species and one outgroup species *Musca domestica* and found that, although predicted tRNA structure-function maps are generally highly conserved in flies, one tRNA Class-Informative Feature (CIF) within the rapidly-evolving ion-binding pocket — Cytosine 17 (C17), ancestrally informative for lysylation identity — independently gained asparaginylation identity and substituted in parallel across tRNA^Asn^ paralogs at least once, possibly multiple times, during evolution of the genus. In *D. melanogaster*, most tRNA^Lys^ and tRNA^Asn^ genes are co-arrayed in one large heterologous gene cluster, suggesting that heterologous gene conversion as well as structural similarities of tRNA-binding interfaces in the closely related asparaginyl-tRNA synthetase (AsnRS) and lysyl-tRNA synthetase (LysRS) proteins may have played a role in these changes. A previously identified Asn-to-Lys anticodon shift substitution in *D. ananassae* may have arisen to compensate for the convergent and parallel gains of C17 in tRNA^Asn^ paralogs in that lineage. Our results underscore the functional and evolutionary relevance of our tRNA structure-function map predictions and illuminate multiple genomic and structural factors contributing to rapid, parallel and compensatory evolution of tRNA multigene families.

## Introduction

Transfer RNAs (tRNAs) were the first family of RNAs to be directly sequenced (Holley 1965) and the first to be solved by X-ray crystallography (Holley *et al*. 1965). Historically, algorithms to estimate phylogeny and substitution patterns were tested on tRNA genes (Cedergren *et al*. 1981; Eigen *et al*. 1989). Early in the genome era, it was reported that tRNA genes can evolve to switch their functional (codon-reading) identities through anticodon shift substitutions, which entail both synonymous and nonsynonymous substitutions in anticodons (Saks *et al*. 1998). However, the small sizes and high similarities of tRNA genes pose obstacles to inferring their orthology, which is needed to better understand the evolutionary processes underlying functional turnover of tRNA genes. An important step forward came from the “micro-syntenic” approach to infer tRNA gene orthology using flanking sequences, first applied in *Drosophila* (Rogers, Bergman, *et al*. 2010) and later to other eukaryotes (Rogers and Griffiths-Jones 2014). These studies revealed that functional turnover of tRNA genes through anticodon shift substitutions is more frequent and widespread than previously known. However, Rogers and Griffiths-Jones (2014) were unable to discern whether anticodon shift substitutions occur more often within or between the two conserved and ancient superfamilies of aminoacyl-tRNA synthetases (aaRSs), called Classes I and II (Eriani *et al*. 1990). The two superfamilies may be further divided into three subclasses each (Cusack 1997), all of which pre-date the divergence of bacteria, archaea and eukaryotes, as exemplified by the consistency with which aaRS paralogs may be used to root the statistical Tree of Life (Brown and Doolittle 1995).

More recent advances in ortholog estimation for tRNA genes exploited positional homology and the organization of tRNA genes as repeated elements in tandem gene arrays, revealing both the great extent of functional turnover and an important role for gene conversion in tRNA evolution (Bermudez-Santana, *et. al*. 2010, Velandia-Huerto *et al*. 2016). These advances in ortholog estimation for tRNA genes have made it possible for the first time to undertake a detailed analysis of substitution rates and patterns in tRNA genes, which is one of two requirements to understand how tRNA genes evolve to switch functions. The second requirement is a means to predict the functional significance of tRNA sequence features — what we call the “tRNA structure-function map.”

In earlier work, we developed an approach to estimate tRNA structure-function maps from pooled, structurally aligned tRNA gene complements inferred from one or more related genomes. Our approach integrates sequence information across all tRNA functional gene families at once, using statistics on structure-conditioned functional information (Freyhult et al. 2006). The relevance of our “information criterion” to predict the tRNA structurefunction map stems from the biophysics of translation, assuming promiscuous interactions across all species of tRNA-binding proteins and tRNAs co-expressed in the same cellular domain or compartment, with association rates that increase proportionally with concentrations (which we estimate for tRNAs by proxy from gene copy numbers, as in the tRNA Adaptation Index, dos Reis *et al*. 2004; Sabi *et al*. 2017) and with aminoacylation probabilities that depend on matching and mis-matching of structural and dynamic (motional) features across all interacting species operating in parallel (Collins-Hed and Ardell 2020). No matter how the phenotypic expression of a given base or base-pair contributes to the classification of tRNAs into substrates and non-substrates by tRNA-binding proteins, whether directly at the binding interface, as a substrate for a critical base modification, or indirectly, by contributing to recognition by base modifying enzymes, through indirect effects on the shape and motion of entire molecules, or via allosteric circuits connecting binding interfaces and active sites in complexes (Sethi *et al*. 2009), the theoretical expectation under selection for translational accuracy is that the genetic bases of those identifying phenotypic traits will become increasingly restricted to the tRNA functional classes that rely on them for their identities (Collins-Hed and Ardell 2020). We call our predictions Class-Informative Features (CIFs) and visualize them in graphs called function logos (Freyhult et al. 2006).

Other authors have more recently applied information statistics to measure the conservation of features within tRNA gene families analyzed independently of one another, in order to predict tRNA identity elements (Tamaki *et al*. 2018, Zamudio and José 2018). The first set of authors combined this information with structural and bioinformatic models of tRNA-protein interaction interfaces. Both sets of authors employ a “conservation criterion” to integrate tRNA sequence variation within functional classes, based on the difference *H*(*X*) – *H*(*X_l_*|*Y* = *y*) between an empirical or assumed composition-based background entropy of structural features *H*(*X*) and the entropy of structural features *H*(*X_l_*|*Y* = *y*) at a site *l* in tRNA genes of a given function *y*. Instead, we employ an “information criterion” that integrates tRNA sequence variation across functional classes, based on the difference *H*(*Y*) – *H*(*Y*|*X_l_* = *x*) between an empirical background entropy of tRNA functions *H*(*Y*) and the entropy of functions *H*(*Y*|*X_l_* = *x*) among tRNA genes that embody structural feature *x* at site *I*. The information criterion has the operational interpretation that tRNA-binding proteins themselves exploit the information derived from CIFs to identify their substrates; while the conservation criterion risks conflating generic tRNA features with class-specific ones, overlooking potential alternative informative feature sets for the same functional identity, and discounting co-evolution of tRNA genes with tRNA-binding protein genes such as aminoacyl-tRNA synthetases. Indeed, using other machine learning approaches, Galili *et al*. (2016) predicted tRNA identity elements that varied across archaeal phyla. In our own work, we have shown that tRNA Class-Informative Features (CIFs) vary across Bacteria (Ardell and Andersson 2006) and we were then able to use this variation to address a challenging problem in the phylogenetics of Alphaproteobacteria (Amrine *et al*. 2014). More recently, we showed that our supervised machine learning phyloclassification algorithm CYANO-MLP could robustly classify cyanobacterial genomes and hindcast the cyanobacterial progenitor of plastids based solely on tRNA CIFs and tRNA gene complements (Lawrence *et al*. 2019).

Even though our tRNA structure-function maps are based on an information criterion rather than a conservation criterion, we recently showed that tRNA CIFs, including Class-Informative Base-Pairs (CIBPs) and Class-Informative Mis-Pairs (CIMPs), are highly conserved within trypanosomes and between trypanosomes and humans, even while showing evidence of co-evolutionary divergence (Kelly *et al*. 2020). Furthermore, our tRNA CIF annotations could predict differential susceptibility to inhibition by chemical agents of homologous aminoacyl-tRNA synthetases (aaRSs) from trypanosomes and humans (Kelly *et al*. 2020). We also found that tRNA gene clusters are conserved in trypanosomes and show clear evidence of rapid evolution by duplication, deletion, and rearrangements (Kelly et al. 2020) consistent with findings from genome comparisons of diverse eukaryotic groups (Bermudez-Santana *et al*. 2010, Velandia-Huerto *et al*. 2016).

In earlier work, as part of the *Drosophila* Twelve Genomes Consortium, we contributed tRNA gene annotations to FlyBase (Bergman and Ardell 2014) and an analysis combining data from multiple ortholog sets of microRNA genes to estimate structurally partitioned evolutionary rates over different structurally-defined categories of sites and site-pairs (*Drosophila* 12 Genomes Consortium 2007). In the present work, we extend our molecular evolutionary analysis approach to *Drosophila* tRNA genes, exploiting their high conservation and structural conformity (Wolfson *et al*. 2001) to estimate and compare evolutionary rates across different tRNA structural components of all functions at individual single- and paired-sites in tRNA genes from twelve species of *Drosophila*. As part of this work, we developed a strategy to optimize the exclusion of species to maximize the length of concatenated alignments across tRNA ortholog sets. After fitting models to our optimized alignments with MrBayes (Huelsenbeck and Ronquist 2001; Ronquist et al. 2012), we discovered that one of several metal ion-binding pockets has evolved rapidly in multiple functional classes of *Drosophila* tRNA genes. Integrating this with tRNA CIF predictions in thirteen species of flies, we found that this rapid evolution is associated with repeated, convergent gains (and possible losses and/or regains) of a tRNA CIF in one tRNA functional gene family, resulting in parallel substitutions in multiple tRNA genes with orthologs on different chromosomes in *D. melanogaster*. We were able to identify multiple structural and genomic factors that have likely contributed to this convergent evolution of tRNA CIFs in the *Drosophila* genus. Our results suggest that anticodon shift substitutions may play a compensatory role in evolution of the tRNA structure-function map.

## Results

We built a custom database of tRNA genes for the *Drosophila* twelve genomes based on FlyBase release 2008-07 (Tweedie et al. 2009) downloaded on October 19, 2011, which contained a total of 3494 tRNA genes. On this set, we integrated orthology annotations from Rogers, Bergman, et al. (2010), COVE scores from tRNAscan-SE 1.3.1 (Lowe and Eddy 1997) and initiator tRNA gene predictions from TFAM v1.3 (Ardell and Andersson 2006, Tåquist et al. 2007). We refolded FlyBase tRNA gene models in tRNAscan-SE and ARAGORN v1.2.34 (Laslett and Canback 2004) as an additional source of functional and structural predictions. Before turning to our main body of results concerning molecular evolution and Class-Informative Features in *Drosophila* tRNA genes, we briefly note an important discovery that we made regarding a previously described anticodon shift substitution in light of our updated annotations of initiator tRNA genes. Our tRNA gene annotations are provided in Supplementary Data.

### *D. simulans* Contains a Non-Canonical Initiator tRNA Gene Created by an Anticodon Shift Substitution

Initiator tRNA genes are a distinct functional class of proteins that read start codons only, and ordinarily carry CAU anticodons in common with elongator tRNA^Met^_CAU_ genes. The first generation of production tRNA gene-finders, tRNAscan-SE v. 1 and Aragorn, erroneously annotate initiator tRNA genes as elongator tRNA^Met^ genes based on predicted anticodons. However, tRNAscan-SE v.2.0 (Chan *et al*. 2019) and TFAM both use sequence profiles to functionally annotate initiator tRNA genes. We found that TFAM could annotate initiator tRNA gene predictions consistently with both experimentally based annotations in *D. melanogaster*, and also across species, in that if TFAM annotated any gene in any ortholog set from Rogers, Bergman, *et al*. 2010 as an initiator tRNA gene, then every other gene in that ortholog set would also be independently annotated by TFAM as an initiator tRNA gene (Bergman and Ardell 2014 and this work). We annotated between five and seven initiator tRNA genes per *Drosophila* genome, for a total of 70 annotated initiator tRNA genes across all species, of which 66 belonged to the ortholog sets from Rogers, Bergman, *et al*. 2010.

We found that one alloacceptor anticodon shift substitution (which converts the functional identity of a tRNA from one amino acid to another), previously reported and validated as tRNA^Met^_CAU_ → tRNA^Thī^_CGU_ (or in their notation, CAT:M → CGT:T) in *Drosophila simulans* (Rogers, Bergman, *et al*. 2010; Rogers and Griffiths-Jones 2014) in fact represents the evolution of a non-canonical initiator tRNA gene in *D. simulans*, in the sense that its predicted tRNA product does not contain the methionine anticodon CAU. This non-canonical initiator tRNA gene in *D. simulans* with FlyBase gene ID gn0256165 and transcript ID tr0296323, evolved as part of ortholog set 183. This ortholog set contains initiator tRNA orthologs in all eight species of Sophophora excluding *D. willistoni*. The *D. simulans* ortholog carries only a single A → G anticodon shift substitution at Sprinzl position 35 (as described in Sprinzl *et al*. 1998, Sprinzl coordinates are a standardized coordinate system for the consensus universal secondary structure of tRNAs as well as conserved, more sub-function-specific structures like the long variable arms of tRNA^Leu^ and tRNA^Ser^). The non-canonical initiator tRNA gene is identical in sequence to all other initiator tRNA genes that we annotated in *D. simulans*, except for the single anticodon shift substitution. Further analysis is required to explore the functional significance of this gene.

### Optimizing Species Exclusion to Maximize Gap-Free Alignment Length of Concatenated tRNA Ortholog Sets

We undertook a global and unbiased analysis of site-specific substitution rates and patterns in *Drosophila* tRNA genes at single-site and site-pair resolution. To do this we analyzed a total of 753 orthologous gene sets from Rogers, Bergman, *et al*. (2010) encompassing 3218 unique FlyBase tRNA transcript IDs, a subset of our reannotated FlyBase set. We found the representation of species to be uneven across the Rogers, Bergman, *et al*. (2010) ortholog sets. Only 47 sets, about 6%, contained orthologs from all 12 *Drosophila* species (Table 1). One species in particular, *D. willistoni*, was represented in very few sets. We proceeded to remove this species from our subsequent substitution rate analyses. We also removed ortholog sets that contained isoacceptor or alloacceptor anticodon shift substitutions, uncertain functional annotations, or predicted pseudogenes.

**Table 1.** Statistics on alignments of concatenated tRNA ortholog sets from *Drosophila* species subsets.

| Species Subset | Species | Ortho. Sets<sup>a</sup> | Sites (gap-free) | Var. Sites (Partitions)<sup>b</sup> | P.I. Sites (Partitions)<sup>c</sup> |
| --- | --- | --- | --- | --- | --- |
| 101111110000<sup>d</sup> | 7 | 147 | 10878 (10598) | 2729 (68) | 27 (15) |
| 110011100111<sup>e</sup> | 8 | 85 | 6290 (6134) | 1589 (67) | 19 (15) |
| 111111110111<sup>f</sup> | 11 | 83 | 6142 (5991) | 1554 (67) | 20 (15) |
| 111111111111 | 12 | 47 | 3478 (3389) | 896 (58) | 16 (12) |

<sup>a</sup> Number of complete tRNA ortholog sets (Rogers, Bergman *et al.* (2010)) from that species subset
<sup>b</sup> Number of variable gap-free sites (or variable site-partitions excluding gap-free sites, of 74) over concatenated alignment
<sup>c</sup> Number of Parsimoniously Informative gap-free sites (or partitions excluding gap-free sites, of 74) over concatenated alignment
<sup>d</sup> Sophophora excluding *D. simulans* and *D. willistoni*
<sup>e</sup> All species excluding *D. sechellia*, *D. yakuba*, *D. persimilis* and *D. willistoni*
<sup>f</sup> All species excluding *D. willistoni*

We wished to integrate data from the largest number of ortholog sets that we could in order to compare substitution rates across sites in an unbiased manner. However, we also wished to avoid having a large number of missing or unidentified orthologs in the concatenated alignments that we made from Rogers, Bergman, *et al*. (2010) ortholog sets, so as to reduce error arising from missing data in our results associated with large blocks of gap characters. We therefore computed statistics over 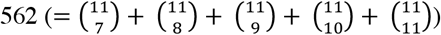 automatically generated concatenated gene alignments corresponding to 562 subsets of genome assemblies from seven or more of the eleven *Drosophila* species excluding *D. willistoni*, in order to find the one species subset that resulted in concatenated alignments of the most ortholog sets and the greatest gap-free length. To make these concatenated alignments for each species subset, first we structurally aligned all tRNA genes together, and then we extracted aligned tRNA gene sequences for every ortholog set containing sequences from at least all the species belonging to the defined species subset. We concatenated these extracted ortholog set alignments together and mapped all sites in this concatenated alignment into separate site-partitions corresponding to each Sprinzl coordinate.

We found that a single species subset, labelled as “101111110000” in Table 1 and representing Sophophora excluding *D. simulans* and *D. willistoni*, yielded the longest gap-free alignment concatenating the most ortholog sets, the greatest numbers of variable and parsimoniously informative sites, and the greatest average number of pairwise differences per site among all alignments we examined. We proceeded to focus on this alignment for downstream analysis as well as two other ones for the sake of comparison: a randomly picked one with a nearly complementary pattern of species exclusion (“110011100111,” excluding *D. sechellia, D. yakuba, D.persimilis* and *D. willistoni*), and one with all but one species (“111111110111,” excluding *D. willistoni*).

### Substitution Patterns in *Drosophila* tRNA Genes

To compute site-specific substitution rates from our concatenated alignments of tRNA ortholog sets, we compared substitution rates per site or per site-pair across different site-partitions of our concatenated alignments corresponding either to tRNA secondary structural elements or to individual sites defined by Sprinzl structurally standardized coordinates (Sprinzl and Vassilenko 2005), using MrBayes 3.2.1 (Huelsenbeck and Ronquist 2001; Ronquist *et al*. 2012), aggregating over different functional classes of tRNA gene orthologs and using the fixed known species tree (*Drosophila* 12 Genomes Consortium 2007). We fitted unpaired sites and loops with the General Time Reversible model (Lanave *et al*. 1984; Tavaré 1986; Rodríguez *et al*. 1990) with invariant sites (GTR+I), and paired-sites and stems with the Doublet(GTR)+I model, also with invariant sites (Ronquist *et al*. 2012). We obtained very similar results fitting the (GTR+I+Gamma) model (data provided in Supplementary Materials) or using the Hasegawa-Kishino-Yano (HKY) model (Hasegawa *et al*. 1985) with or without Gamma-distributed site-rate variation (Phillips 2003 and Figures S1 and S2 in Supplementary Online Materials). We ran MrBayes with the option “ratemult=scaled,” which implies that all rates reported and shown in Figures 1 and 2 are scaled so that the mean rate across all site-partitions is 1.0 substitutions per site or site-pair.

**Figure 1.**
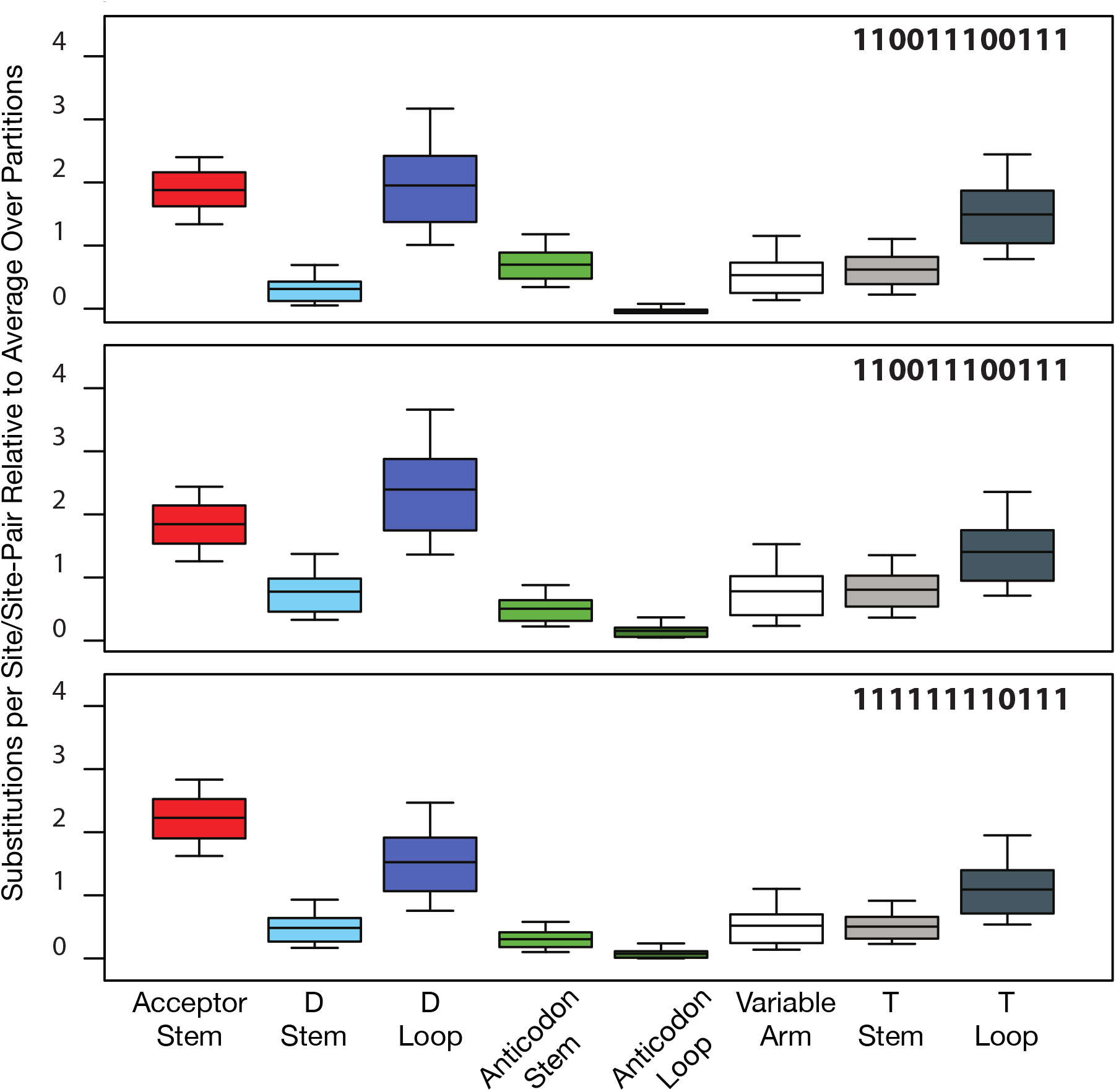
95% Credible intervals, interquartile ranges and medians of relative substitution rates by secondary structural elements in *Drosophila* tRNA genes as calculated in MrBayes 3.2.1 using the GTR+I/Doublet(GTR+I) models for loops/stems and the partitioned concatenated alignments indicated by bit-sets as defined in Table 1.

**Figure 2.**
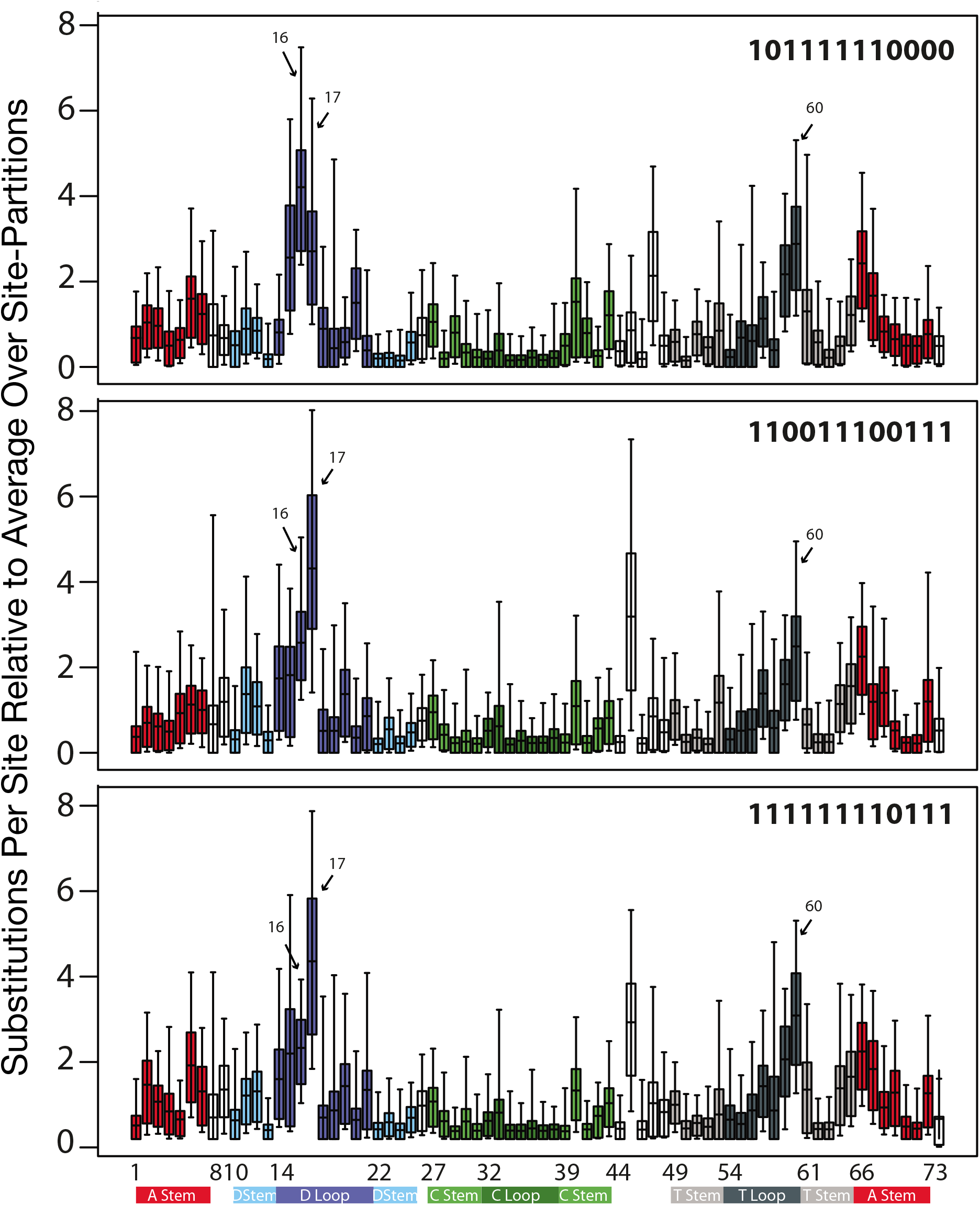
95% Credible intervals, interquartile ranges and medians of relative substitution rates by individual site/Sprinzl coordinate in *Drosophila* tRNA genes as calculated in MrBayes 3.2.1 using the GTR+I model for individual sites and the partitioned concatenated alignments indicated by bit-sets as defined in Table 1.

In our coarse-grained analysis of substitution rates by secondary structural elements we observed that, regardless of which alignments we used, the largest substitution rates occur in the D-loop and acceptor stem followed closely by the T-loop (Figure 1). Considering the relative constraint that acceptor stems are expected to have from identity-driven interactions with proteins (Giegé, Sissler, *et al*. 1998), the acceptor stem shows a surprisingly high rate of evolution. Furthermore, regardless of which alignments or models we used, in our analysis of substitution rates by individual sites, we found that three sites — corresponding to Sprinzl coordinates 16, 17, and 60 — showed markedly higher rates of substitution than any other, followed by sites 15, 45 and 59 (Figure 2).

Sprinzl coordinates 16, 17 and 60 are three among eight sites that form an extended ion-binding pocket (Sprinzl coordinates 15, 16, 17, 18, 19, 20, 59, and 60) between the D- and T-arms at the interface of the two tRNA helical domains (Behlen *et al*. 1990), as shown in Figure 3. This pocket was first described as site 1 in the original orthorhombic crystal form (Holbrook et al. 1977) and site 3 in the monoclinic structure (Jack *et al*. 1977) and was later shown to potentially bind different metal ions including magnesium, cobalt, manganese, and lead (Jack *et al*. 1977; Holbrook *et al*. 1977; Behlen *et al*. 1990; Shi and Moore 2000). More recent structural work better resolved the structure of Sprinzl site 16, which is a fairly conserved uracil among eukaryotes (Marck and Grosjean, 2002), most likely post-translationally modified to dihydrouridine, and demonstrated the conformational sensitivity of this to ionic conditions (Shi and Moore 2000). The hypergeometric probability that three sites with elevated substitution rates occur within the eight sites that form the extended ion-binding pocket, out of 74 possible sites, is 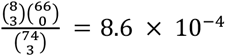.

**Figure 3.**
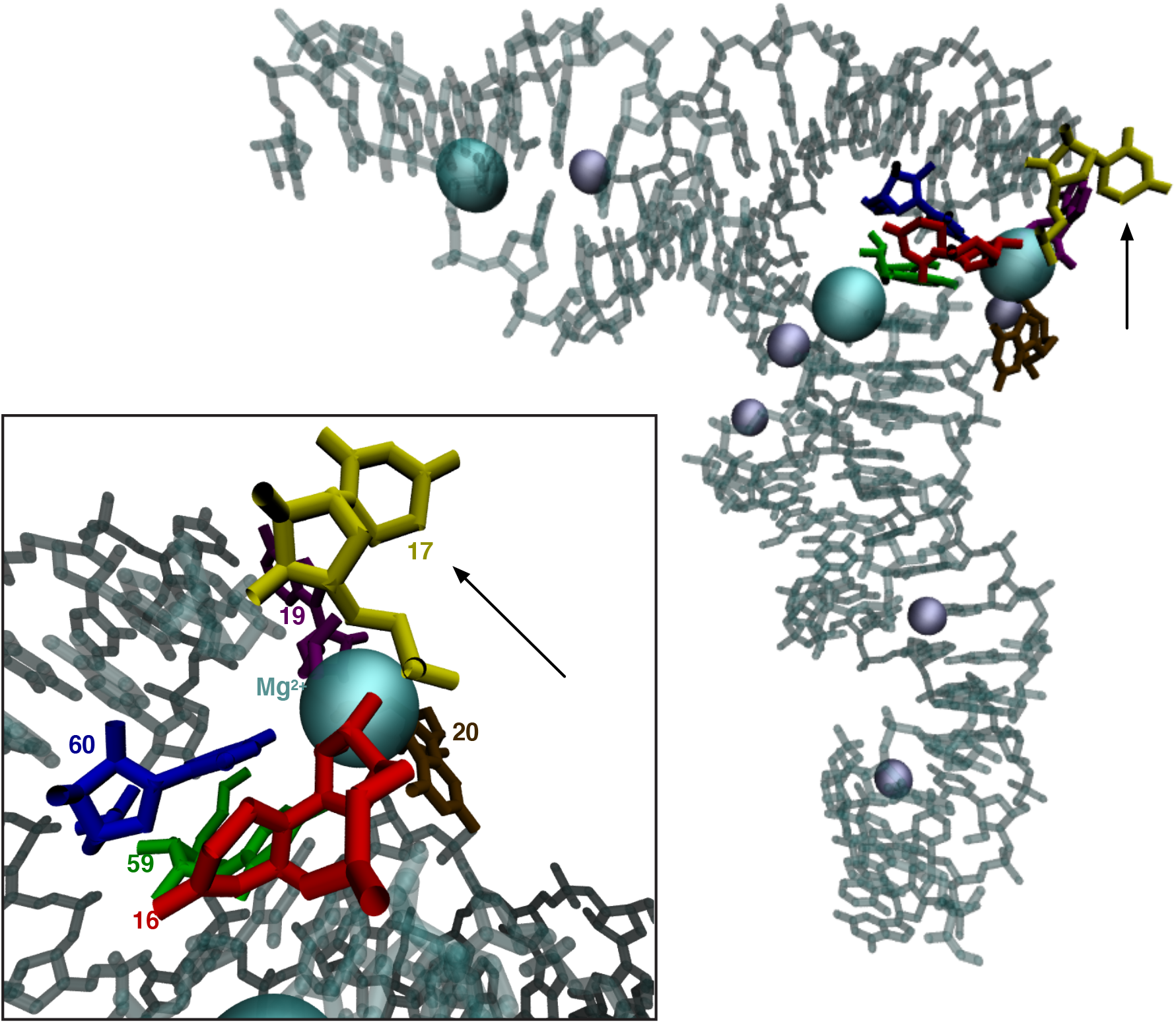
Rapidly-evolving sites in *Drosophila* tRNAs are part of a conserved, structurally plastic ion-binding pocket near the interface of the two tRNA helical domains, indicated by colored nucleotides and labelled in the inset panel at higher magnification. This image visualizes PDB structure accession ID 1EHZ (Shi and Moore 2000) and rendered in VMD 1.9.2 (Humphrey et al. 1996).

In the alignment with the most species (111111110111), we found that 45 ortholog groups spanning 14 functional classes carry substitutions in sites 16, 17 and 60 of this ion-binding pocket, as shown in Table S1 of Supplementary Materials.

Chromosomal gene location was significantly associated with substitutions in pocket sites 16, 17 and 60 (*X*^2^ = 56.3266, *N* = 692, *p* = 6.96 × 10”^11^), and extended pocket sites 15, 16, 17, 18, 19, 20, 59, and 60 (*X*^2^ = 87.4858, *N* = 726, *p* < 2.2 × 10^-16^), Supplementary Materials, Table S2. Substitutions in sites 16, 17 or 60 of the pocket were not associated with alloacceptor class switches overall (*X*^2^ = 20.9616, *N* = 672, *p* = 0.523), nor were substitutions in the extended pocket sites 15, 16, 17, 18, 19, 20, 59, and 60 (*X*^2^ = 23.2573, *N* = 703, *p* = 0.387). Ortholog sets containing anticodon shift substitutions more often exhibited substitutions in pocket sites 16,17, and 60 (*X*^2^ = 8.8797, *N* = 713, *p* = 0.00288), but not extended pocket sites 15, 16, 17, 18, 19, 20, 59, and 60 (*X*^2^ = 2.7304, *N* = 696, *p* = 0.0985). We provide estimates of transition and transversion rates for *Drosophila* tRNA genes in Supplementary Materials Table S3.

### Class-Informative Base-Pairs and Base-Mispairs in *Drosophila melanogaster* tRNAs

As described in Kelly *et al*. (2020), we reimplemented our previously published algorithm to estimate structure-conditioned functional information statistics from Freyhult *et al*. (2006) extending it to compute functional information for all sixteen possible base-pair or base mis-pair features occurring in any paired sites of the consensus secondary structure of tRNAs. The program accepts as input a set of multiple alignments, one for each functional sub-family of any RNA or protein multigene family, all mutually structurally aligned, and computes as its output function logo visualizations and tables of statistics on CIFs and their evolution in one or more taxa. The software, called tSFM (“tRNA Structure-Function Mapper”) v1.0.1 despite being applicable to any RNA or protein family, uses a permutation approach to measure the significance of CIFs and CIF evolution dissimilarities and corrects for multiple comparisons by controlling Family-Wise Error Rates (FWERs) or False Discovery Rates, such as that of Benjamini and Hochberg (1995). tSFM v.1.0.1 and later versions are freely available for download on GitHub at https://github.com/tlawrence3/tSFM.

We computed tRNA Class-Informative Base-Pairs and Mis-Pairs from structurally aligned *Drosophila melanogaster* tRNA genes filtered from our custom gene annotation, with tRNAscan-SE 1.3.1 COVE scores of at least 50 bits (tRNAscan-SE 1.3.1 COVE scores estimate the log-likelihood that a given sequence conforms to the consensus primary and secondary structure of tRNAs in general, as opposed to a random sequence of the same composition), removing tRNA genes of indeterminate function and selenocysteine tRNA genes, and leaving 288 tRNA genes remaining. We then computed tRNA Class-Informative Base Pairs and Base Mis-Pairs from these data using tSFM v.1.0.1, and only retained features with a Benjamini-Hochberg False Discovery Rate of 5%, as shown in Figure 4. The total height of a stack of letters at any site quantifies the information potentially gained about the functional type of a tRNA by a tRNA-binding protein if it recognizes the specific type of paired-site feature corresponding to that location. The letters within each stack symbolize functional types of tRNAs, wherein IUPAC one-letter amino acid codes represent elongator tRNA aminoacylation identities and “X” symbolizes initiator tRNAs. The relative heights of letters within each stack quantifies the over-representation of tRNA functional types carrying that feature relative to the background frequency determined by gene frequencies of functional types (as calculated through the normalized log-odds). Thus, for example a U1:A72 base-pair at Sprinzl coordinates 1 and 72 (or some modification that biosynthetically depends on that base-pair) is informative for tRNA^Asp^ and tRNA^Glu^ in *D. melanogaster*.

**Figure 4.**
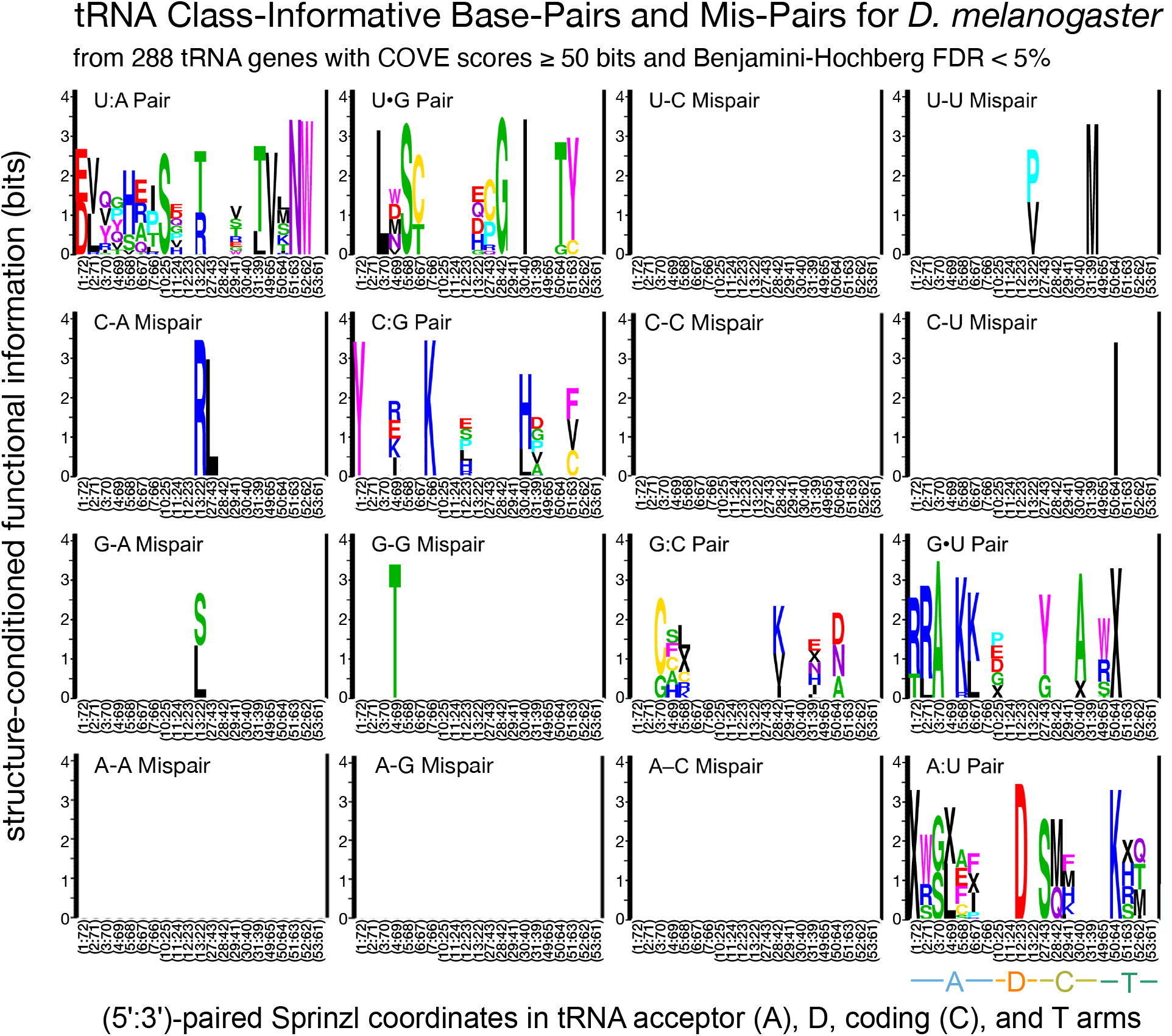
Class-Informative Base-Pairs and Base Mis-Pairs from 288 reannotated tRNA genes in *Drosophila melanogaster* with COVE scores of at least 50 bits as computed in tSFM v1.0.1 with a Benjamini-Hochberg False Discovery Rate of 5% (CIFs not meeting this significance threshhold are not shown). The total height of a stack of letters quantifies information gained about the functional type of a tRNA by a tRNA-binding protein if it specifically recognizes the paired features indicated. The letters within each stack symbolize functional types of tRNA genes, wherein IUPAC one-letter amino acid codes represent elongator tRNA aminoacylation identities and “X” symbolizes initiator tRNAs. The relative heights of letters within each stack quantifies the over-representation of tRNA functional types carrying that feature relative to the background frequency determined by the frequencies of genes of various functional types (as calculated through normalized log-odds).

Figure 4 shows that a great deal of functional information resides in Base-Pairs and Base Mis-Pairs of *Drosophila melanogaster* tRNAs. Several base-mispairs are functionally informative, including U13:U22 in the D-arm, which is associated to both tRNA^Pro^ and tRNA^Val^ (contained in 32 tRNA genes of 32 total), U31:U39 in the C-arm contained in all six tRNA^Met^ genes, C50:U64 in the T-arm contained in nine of 11 tRNA^Ile^ genes, G13:A22 in the D-arm contained in all 42 tRNA^Ser^ and tRNA^Leu^ genes, and a G4:G69 mis-pair in the acceptor stem associated with nine of 16 tRNA^Thr^ genes. We show a plot of the significance of paired-site CIFs as a function of CIF information in Supplementary Figure S3. A text-file containing all statistics for paired-site CIFs in *D. melanogaster*, including frequencies of CIFs in genes of various functions, as well as *p*-values and FDRs, is provided in the Supplementary Code and Data.

### Conservation of Class-Informative Features in *Drosophila* tRNA Genes

The wealth of information we have on *Drosophila* evolution provides an unprecedented opportunity to understand how tRNA Class-Informative Features evolve. To examine the conservation of tRNA CIFs in *Drosophila*, we computed tRNA CIFs in each species independently in the same way as we did for *D. melanogaster*. In order to polarize evolutionary changes in tRNA CIFs we additionally analyzed tRNA gene annotations and CIFs in the outgroup species *Musca domestica*, which diverged from *Drosophila* between 20–80 million years ago (Wiegmann *et al*. 2003). As shown in Figure 5, tRNA single-site CIFs are very highly conserved in *Drosophila* and flies generally. Although this figure is designed to emphasize the similarity of CIF estimates across species, separate detailed logo figures for each species as well as combined logos for base-paired features are provided in the Supplementary Online Materials and confirm strong conservation of both Class-Informative Base-Pairs and Class Informative base Mis-Pairs in the *Drosophila* genus. Broadly speaking, our predictions are very nearly identical in all fly genomes we analyzed, with one visibly clear exception standing out: C17, which occurs within the metal ion-binding pocket that we previously identified as having elevated substitution rates.

**Figure 5.**
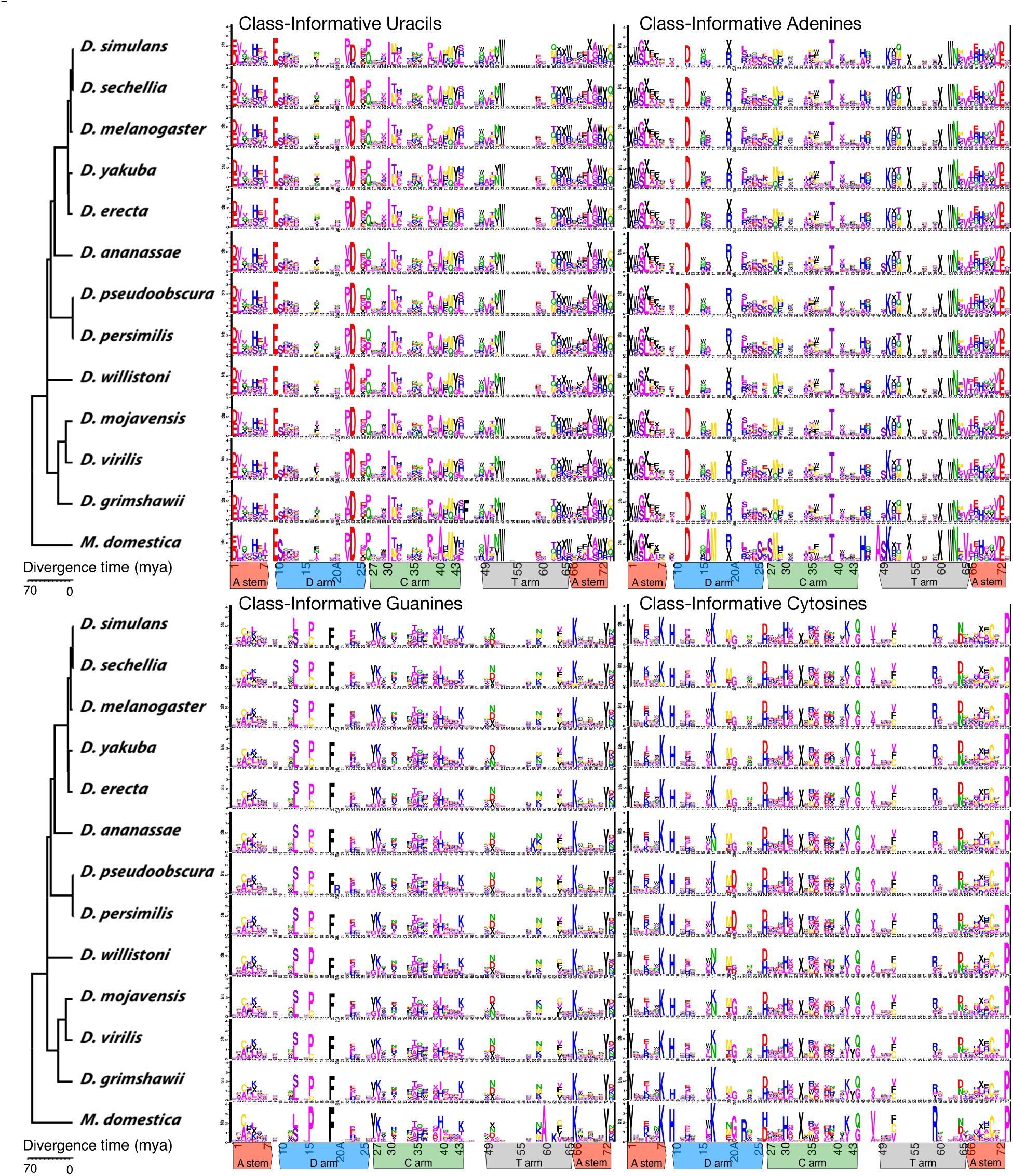
Conservation of tRNA single-site CIFs in *Drosophila* with *Musca domestica* as an outgroup showing extensive conservation of tRNA CIFs in *Drosophila*. Tree topology from *Drosophila* 12 Genomes Consortium (2007), with divergence dates from Tamura et al. (2003) and Hennig et al. (1981).

As shown in Fig. 6, CIF C17 is ancestrally associated with tRNA^Lys^ in flies (as judged by parsikmony from outgroup polarization) but it became associated with all or nearly all tRNA^Asn^ paralogs in at least three phylogenetically distinct *Drosophila* lineages, one of which, leading to *D. ananassae*, also underwent the Asn-to-Lys anticodon shift previously reported in Rogers, Bergman, *et al*. (2010). One interpretation of the evolution of site 17 in tRNA^Asn^ genes is that of multiple gains of C17, but losses and regains cannot be excluded. To underscore the validity and biological significance of our CIF evolution results, we emphasize that all of the 138 tRNA^Asn^ genes analyzed in Fig. 6 have COVE scores of 50 bits are greater — meaning they are above a typical threshold used for inclusion in tRNA gene annotation gene-sets — and in fact, all but two of them have scores above 70 bits — consistent with well-folding, typical tRNA sequences — while all but six have scores above 80 bits (for context, the maximum COVE bit-score over our entire annotation set is 87.38 bits). In addition, 86 of 111 FlyBase tRNA^Asn^ genes belong to the Rogers *et al*. ortholog sets. For comparison’s sake, ten of 111 FlyBase records for tRNA^Asn^ genes have COVE scores below 50 bits, of which two belong to the Rogers et al. ortholog sets. Of the 29 genes that substituted to C17 that belong to the Rogers et al. ortholog sets, 15 are considered “core genes” in the sense of Rogers, Bergman, *et al*. 2010. All of them have predicted Asn anticodons and were annotated as tRNA^Asn^ genes by TFAM. In Fig. 7, we show the raw alignment data for tRNA^Asn^ and tRNA^Lys^ genes in *D. ananassae* and tRNA^Asn^ genes in *D. melanogaster*, demonstrating their extreme consistency in sequence with very little variation outside site 17 and the anticodon shift substitution in *D. ananassae*. While our interpretation rests on the assumption that we have correctly annotated the function of these tRNA^Asn^ genes and that their functions as such have been conserved during their evolution, overall, we believe that our evidence is strong that tRNA^Asn^ genes that have undergone repeated parallel evolution in *Drosophila*.

**Figure 6.**
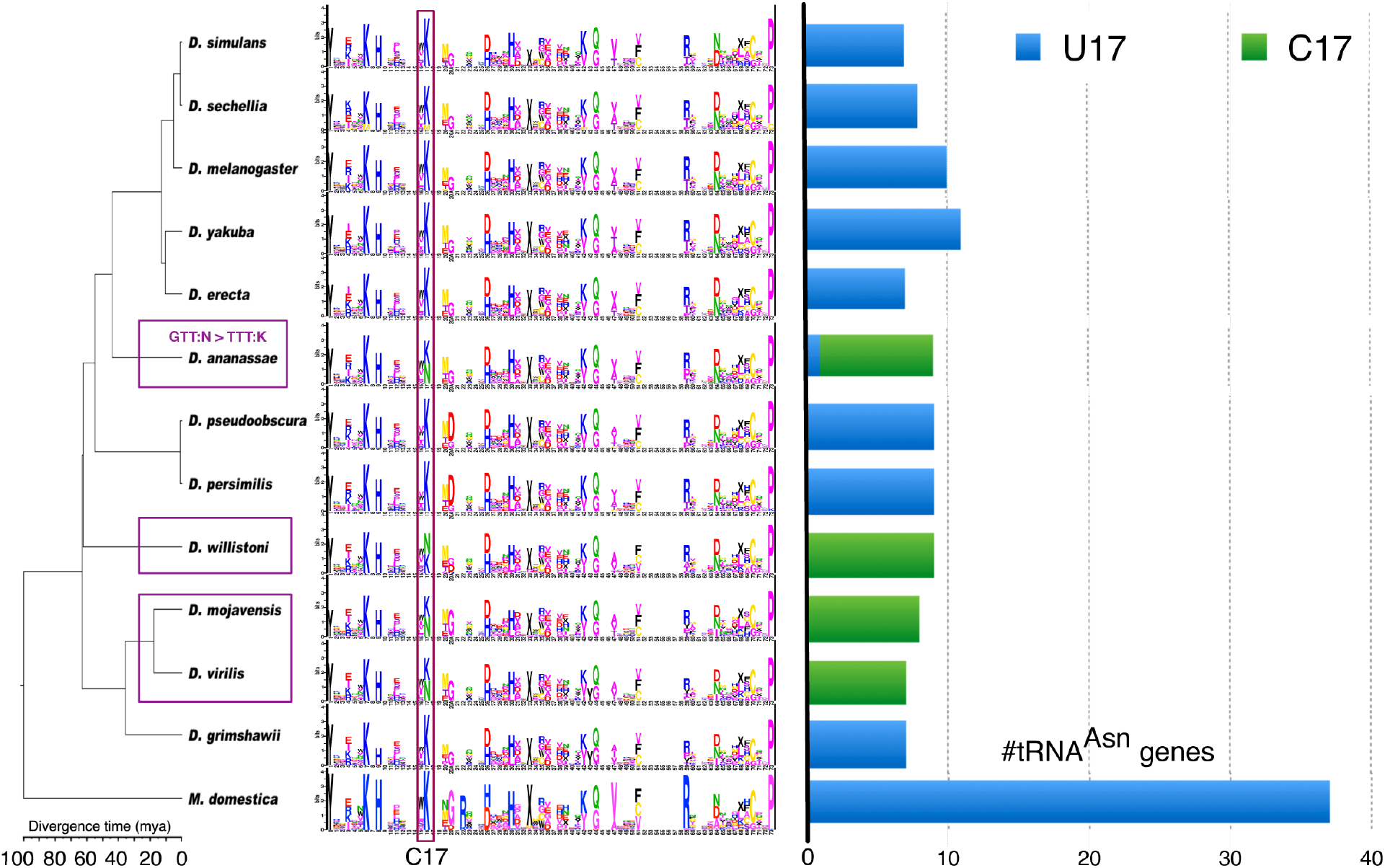
Evolution of functional association of CIF C17 in flies. Cytosine single-site function logos are from Fig. 5, the tree topology is from *Drosophila* 12 Genomes Consortium (2007), and divergence dates are from Tamura *et al*. (2003) and Hennig *et al*. (1981). The three clades in which C17 became associated with tRNA^Asn^ genes are indicated by purple boxes on the tree, one of which (*D. ananassae*), also underwent the Asn-to-Lys anticodon shift previously reported in Rogers, Bergman, et al. (2010) as indicated in top purple box at figure left. The stacked bar graph at figure right shows the frequencies of U17 or C17 across all tRNA^Asn^ genes in each genome showing that C17 was gained and/or lost in parallel and regained across all or mostly all tRNA^Asn^ genes at least three times during evolution of the *Drosophila* genus.

**Figure 7.**
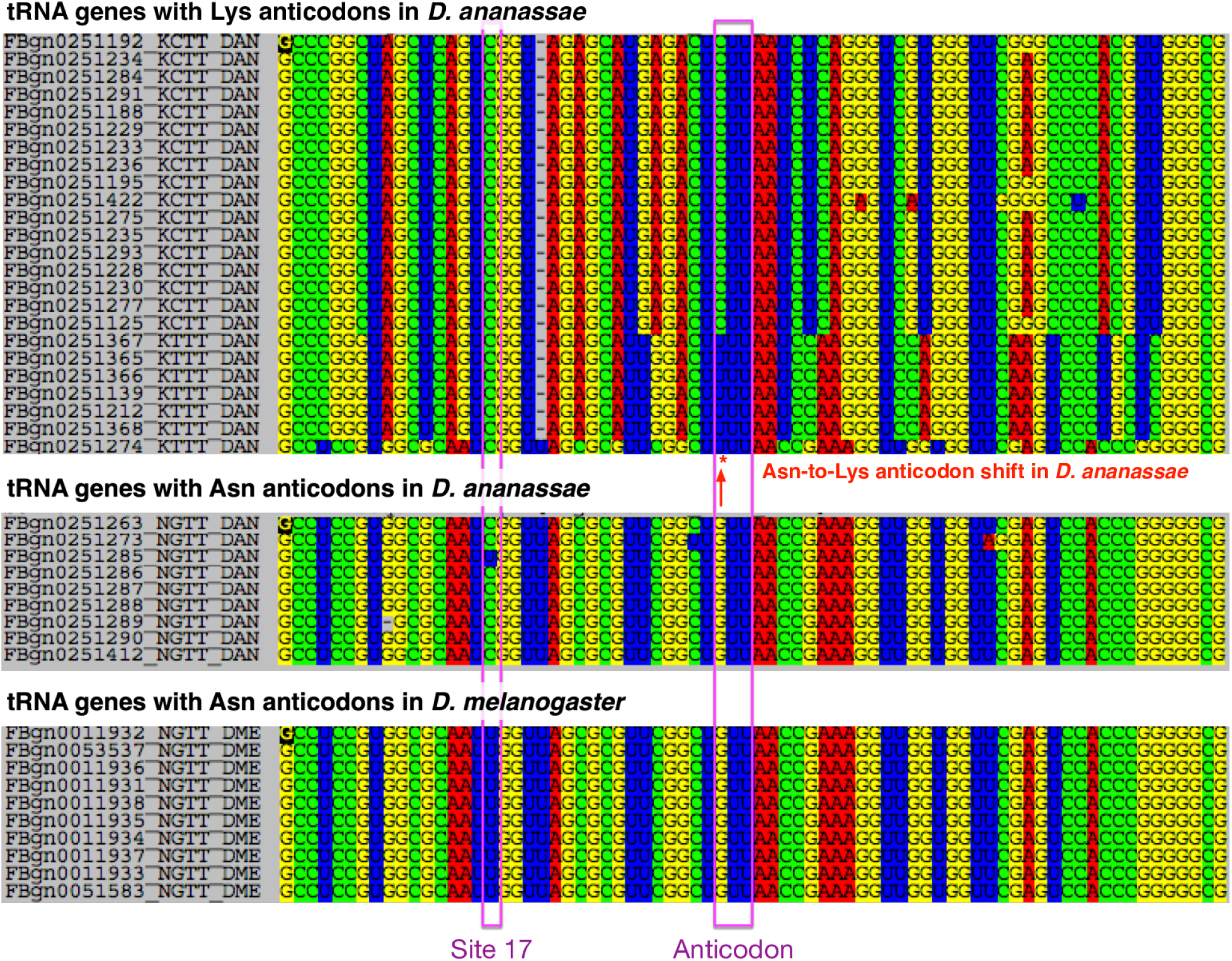
Structural alignments, visualized in SEAVIEW (Gouy *et al*. 2010) of all annotated tRNA^Lys^ and tRNA^Asn^ genes in *D. ananassae* and all annotated tRNA^Asn^ genes in *D. melanogaster*, highlighting sites corresponding to anticodons and site 17, as well as the Asn-to-Lys anticodon shift substitution gene studied in Rogers, Bergman, et al. (2010).

In Figure S4, also in Supplementary Materials, we show that the relative average functional information carried by nucleobases is inversely related to their compositional frequency in *D. melanogaster* tRNA genes (even though the measure does not depend on the compositional entropy of bases themselves). This means that rare bases tend to be more informative for function in *Drosophila* tRNAs. The relative frequency of bases in *D. melanogaster* tRNA genes decreases from G to C to U to A. Their functional information (averaged over all sites and functions) in *D. melanogaster* tRNA CIFs increases correspondingly.

## Discussion

In their analysis of tRNA gene ortholog sets from flies, primates and other groups, Rogers and Griffiths-Jones (2014) reported no evidence of preferential anticodon shift substitutions within or between genes for functional classes of tRNAs charged by either of the two aminoacyl-tRNA synthetase superfamilies, Class I or Class II. However, they did report that an Asn-to-Lys alloacceptor anticodon shift substitution was one of only two types of anticodon shift substitutions that occurred more than once out of a total of 30 types in their dataset, once in *Drosophila* and twice independently in primates. In this work, we show that in flies, the previously reported Asn-to-Lys anticodon shift substitution co-occurs in one lineage with recruitment of CIF C17 to tRNA^Asn^ genes from tRNA^Lys^ genes where it is ancestral, that this co-option of CIF C17 from tRNA^Lys^ to tRNA^Asn^ was multiply gained and/or lost and regained at least three times in *Drosophila* evolution, and that when it was gained (and/or lost and regained), it changed through parallel substitutions across all or mostly all tRNA^Asn^ genes. We note that AsnRS and LysRS are both Class IIb synthetases, suggesting that perhaps co-option of tRNA CIFs occurs more frequently within aaRS sub-classes, perhaps because aaRSs of the same sub-class have more similar binding interfaces on tRNAs (Giegé, Touzé, *et al*. 2008). In our recently published theory for the evolution of aaRS-tRNA interaction networks, tRNA mutations potentially influence interactions with multiple aaRSs when they occur inside shared interfaces (Collins-Hed and Ardell 2019). The pattern we report here is consistent with an interpretation that the Asn-to-Lys anticodon shift substitution evolved to compensate the co-option of C17 from Lys to Asn in the *D. ananassae* lineage. Of course, our interpretations rest generally on the correctness of our functional and structural annotations and the genome assembly and sequence data that underlie them.

Comparing the genomic locations of tRNA^Lys^ and tRNA^Asn^ genes from the FlyBase data and visualizing them in the UCSC Genome Browser for *Drosophila melanogaster*, we found that eight of ten tRNA^Asn^ genes co-occur in a heterologous gene array along with eight tRNA^Lys^ genes in both direct and inverted orientations within approximately 45 Kb on Chromosome 2R, far away from two additional tRNA^Asn^ gene singletons, one on 2R and another on 3R. A detailed look at our integrated annotations (provided in the Supplementary Materials) reveals that ortholog sets of tRNA^Asn^ genes that underwent evolution in site 17 contain *D. melanogaster* genes both inside and outside of this heterologous gene array. Specifically, the Rogers, Bergman *et al*. (2010) ortholog sets of tRNA^Asn^ genes that intersect the *D. melanogaster* heterologous gene array are: sets 46 and 61, containing genes from twelve (respectively eleven) species, including from all four species whose genes underwent substitution to C17; ortholog set 62 with genes from seven species including *D. ananassae, D. willistoni*, and *D. mojavensis* with substitutions to C17; ortholog set 47 with genes from nine species including *D. ananassae, D. willistoni*, and *D. virilis* with C17; ortholog set 48 with genes from seven species including *D. ananassae* and *D. willistoni*, with C17; ortholog sets 50 and 52 which contain five (respectively three) genes including one from *D. ananassae* with C17; and ortholog set 51 with four genes, one from *D. ananassae* with U17. Outside of the heterologous gene array, ortholog set 138, whose *D. melanogaster* ortholog resides elsewhere on Chromosome 2R, contains eleven genes with three genes from *D. ananassae, D. willistoni*, and *D. virilis* that substituted to C17, while ortholog set 205, whose *D. melanogaster* ortholog resides on Chromosome 3R, contains genes from all twelve species, including from all four species whose genes underwent substitution to C17.

While a renewed tRNA gene reannotation effort with long-read assembly data is needed to better understand the causes of the changes we have reported here, the genome location data in *D. melanogaster* is consistent with a scenario in which gene conversion by illegitimate recombination may have played a role in the convergent and parallel evolution of the tRNA structure-function map in *Drosophila*. Furthermore, if the organization of tRNA^Asn^ genes in *D. melanogaster* are broadly representative across species, a simple scenario of concerted gene evolution (Nei and Rooney 2005) is insufficient to explain the parallel and repeated substitutions of C17 (and possibly also U17) in tRNA^Asn^ genes that we observed. Rapid evolution of tRNA gene arrays through rearrangements appears to be universal in eukaryotes (Velandia-Huerto *et al*. 2016) and at least some prokaryotes (Tremblay-Savard *et al*. 2015).

In summary, interactions across at least three different levels of biological structure appear to have contributed to a specific and recurring pattern of rapid, parallel evolution and functional turnover of tRNA genes in *Drosophila*: of physically interacting sites within tRNA tertiary structure, of co-clustered tRNA genes within genomes, and of overlapping interfaces for structurally similar and closely related tRNA-binding proteins. Further work is needed to generalize the observations we have made and to discern their causes.

## Methods

### Data, Ortholog Sets and Alignments

We obtained tRNA gene sequences and annotations for 12 species of *Drosophila* from FlyBase (2008_07 release, McQuilton *et al*. 2012) on October 16, 2011. For further functional re-annotation, we re-folded and re-annotated these tRNA sequences using tRNAscan-SE 1.3.1 (Lowe and Eddy 1997) and ARAGORN 1.2.34 (Laslett and Canback, 2004). The genome for *Musca domestica* (Scott *et al*. 2014) was downloaded from NBCI on October 8th, 2013. The genome was annotated using predictions from tRNAscan-SE 1.3.1 (Lowe and Eddy 1997).

We identified Initiator tRNA genes using TFAM 1.3 (Tåquist *et al*. 2007). We aligned all tRNAs using Infernal 1.1 (Nawrocki *et al*. 2009) with the RFAM covariance model for tRNAs (RF00005) (Burge *et al*. 2012). We edited alignments manually using Seaview 4.3.4 (Gouy et al. 2010) to produce a final alignment of length 74, and mapped each site manually to Sprinzl coordinates (Sprinzl and Vassilenko 2005). We verified our coordinate mapping using tRNAdb (Jühling *et al*. 2009). We retained Sprinzl coordinate 20A and removed the majority of the variable arm, except Sprinzl coordinates 45 through 49, for subsequent analysis.

We downloaded ortholog sets of tRNA genes on October 18, 2011 from supplementary data available at http://gbe.oxfordjournals.org/content/2/467/suppl/DC1 (Rogers, Bergman, *et al*. 2010). To optimize ortholog sets, we calculated lengths of pruned species trees using the Bio::TreeIO module in BioPerl 1.4.0 (Stajich, Block, *et al*. 2002) and calculated numbers of variable and parsimoniously informative sites of alignments, as well as average fraction of pairwise differences, using the Bio::PopGen modules (Stajich and Hahn 2005) in BioPerl 1.4.0.

### Analysis of Substitution Rates

All subsets of data were curated into concatenated alignments by previously published ortholog sets (Rogers, Bergman, *et al*. 2010). Ortholog sets with anticodon shift substitutions, genes of indeterminate function, or pseudogenes were removed from all analyses of substitution rates.

We estimated substitution rates with MrBayes 3.2.1 (Huelsenbeck and Ronquist 2001; Ronquist et al. 2012) using the fixed known species tree (Drosophila 12 Genomes Consortium 2007). For all runs we constrained change to the tree topology by setting rates of stochastic TBR and branch multipliers to zero probability. All Bayesian analyses were run with two simultaneous chains for 4 × 10^6^ iterations, monitoring convergence of split frequency standard deviations and saved parameters every 500 iterations, and with option “ratemult=scaled.”

We computed evolutionary rates by assigning sites or site-pairs into corresponding data partitions. For structural data partitions, we considered nine structural tRNA components: acceptor stem (Sprinzl sites in *Drosophila*: 1 – 7, 67 – 73), D stem (10 – 13, 23 – 26), D loop (14 – 22), anticodon stem (28 – 32, 40 – 44), anticodon loop (33 – 39), variable arm (45 – 49), T stem (50 – 54, 62 – 66), T loop (55 – 61), and “other sites” (8, 9, 27, 74). Sites in “other” are not involved in base-pairs or considered part of loop structures in tRNA. We estimated rates using the General Time Reversible (GTR+I) substitution model (Lanave *et al*. 1984; Tavaré 1986; Rodríguez *et al*. 1990) with Invariant sites and the Doublet(GTR)+I model with Invariant sites (Ronquist *et al*. 2012), allowing stationary state frequencies and all other substitution model parameters to be independent across site-partitions. Substitution rate multipliers for partitions were scaled to have an average rate of one substitution per site/site-pair over all partitions. Alignment data and MrBayes initialization scripts in NEXUS format with corresponding alignments are provided in supplementary data. The first 25% of parameter calculations were discarded as burn-in for statistical analysis. Parameter posterior probabilities were imported to R (R Core Team 2013). We used the coda package (Plummer et al. 2006) for Markov Chain Monte Carlo simulations diagnostics and the lattice package (Sarkar 2008) for multivariate analysis. All data from MCMC simulations is presented with 95% Bayesian credible intervals.

### Detection of Class-Informative Features

For analysis of tRNA CIFs, we removed genes with anticodon shift substitutions, genes of indeterminate function, and pseudogenes and we further filtered genes to have tRNAscan-SE COVE scores (Lowe and Eddy 1997) of at least 50 bits. We identified Class-Informative features (CIFs) for 12 species of *Drosophila* using tSFM v1.0, available from https://github.com/tlawrence3/tSFM, using the Nemenman-Shafee-Bialek (NSB) Bayesian entropy estimator (Nemenman et al. 2002) for features in two or more sequences and an exact estimator (Schneider et al. 1986) otherwise. To compute the significance of paired-site CIFs in *D. melanogaster*, we computed permutation *p*-values (permuting functional class assignments over sequences that contain a CIF) for the total information of paired-sites only, and then computed Benjamini-Hochberg False Discovery Rates (Benjamini and Hochberg 1995) from those *p*-values.

## Supporting information

SOM for Phillips and Ardell 2021

## Data Access

All code and data required to produce results in this work have been deposited to FigShare and freely available at https://doi.org/10.6084/m9.figshare.12713705.v1.

## Acknowledgments

The authors thank Carolin Frank, Michael Cleary, Michael Colvin, Josh Phillips, David Pollock, David Liberles, and three reviewers for helpful comments and discussions. DHA was supported by a National Science Foundation grant (INSPIRE-1344279), a National Institutes of Health grant (AI127582), and a Julius Kuhn Guest Professorship at Martin Luther University in Halle-Wittenberg, United States. Computational research in this report was performed on the MERCED HPC cluster supported by the National Science Foundation, United States (ACI-1429783).

## Declarations

### Ethics approval

Not applicable.

### Consent to participate

Not applicable.

### Consent for publication

All authors consent to publication.

### Authors’ contributions

All authors contributed to the study conception and design. Material preparation, data collection and analysis were performed by Julie Baker Phillips and David Ardell. The first draft of the manuscript was written by Julie Baker Phillips and David Ardell. All authors read and approved the final manuscript.

## References

Amrine KCH, Swingley WD, and Ardell DH. 2014. tRNA signatures reveal a polyphyletic origin of SAR11 strains among alphaproteobacteria. PLoS Computational Biology. 10: e1003454.

Ardell, DH, and Andersson, SGE. 2006. TFAM detects co-evolution of tRNA identity rules with lateral transfer of histidyl-tRNA synthetase. Nucleic Acids Research, 34: 893–904.

Bailly M, Giannouli S, Blaise M, Stathopoulos C, Kern D, and Becker HD. 2006. A single tRNA base-pair mediates bacterial tRNA-dependent biosynthesis of asparagine. Nucleic Acids Res. 34: 6083–6094.

Behlen LS, Sampson JR, DiRenzo AB, and Uhlenbeck OC. 1990. Lead-catalyzed cleavage of yeast tRNA^Phe^ mutants. Biochemistry. 29: 2515–2523.

Benjamini Y and Hochberg Y. 1995. Controlling the false discovery rate: A practical and powerful approach to multiple testing. Journal of the Royal Statistical Society: Series B (Methodological). 57: 289–300.

Bergman, C and Ardell, DH. 2014. Methods for nuclear tRNA gene predictions for 12 species in the genus *Drosophila*. FigShare. Journal contribution. https://doi.org/10.6084/m9.figshare.1233438.v1

Bermudez-Santana, C., Attolini, C.SO., Kirsten, T. et al. (2010). Genomic organization of eukaryotic tRNAs. BMC Genomics 11:270.

Brown JR and Doolittle WF. 1995. Root of the universal tree of life based on ancient aminoacyl-tRNA synthetase gene duplications. Proc Natl Acad Sci U S A. 92: 2441–2445.

Burge SW, Daub J, Eberhardt R, Tate J, Barquist L, Nawrocki EP, Eddy SR, Gardner PP, and Bateman A. 2012. Rfam 11.0: 10 years of RNA families. Nucleic Acids Research. 41: D226–D232.

Cedergren RJ, Sankoff D, LaRue B, and Grosjean H. 1981. The evolving tRNA molecule. CRC Critical Reviews in Biochemistry. 11: 35–104.

Chan PP, Lin BY, Mak AJ, Lowe TM. 2019. tRNAscan-SE 2.0: Improved detection and functional classification of transfer RNA genes. bioRxiv 614032; doi: https://doi.org/10.1101/614032.

Collins-Hed AI and Ardell DH. 2019. Match fitness landscapes for macromolecular interaction networks: Selection for translational accuracy and rate can displace tRNA-binding interfaces of non-cognate aminoacyl-tRNA synthetases. Theor Popul Biol. 129: 68–80.

Cusack S. 1997. Aminoacyl-tRNA synthetases. Current Opinion in Structural Biology. 7: 881–889.

dos Reis, M., Savva, R., & Wernisch, L. 2004. Solving the riddle of codon usage preferences: a test for translational selection. Nucleic Acids Research, 32: 5036–5044.

*Drosophila* 12 Genomes Consortium. 2007. Evolution of genes and genomes on the *Drosophila* phylogeny. Nature. 450: 203–18.

Eigen M, Lindemann BF, Tietze M, Winkler-Oswatitsch R, Dress A, and von Haeseler A. 1989. How old is the genetic code? Statistical geometry of tRNA provides an answer. Science. 244: 673–679.

Eriani G, Delarue M, Poch O, Gangloff J, and Moras D. 1990. Partition of tRNA synthetases into two classes based on mutually exclusive sets of sequence motifs. Nature. 347: 203–206.

Freyhult E, Moulton V, and Ardell DH. 2006. Visualizing bacterial tRNA identity determinants and antideterminants using function logos and inverse function logos. Nucleic Acids Research. 34: 905–916.

Galili T, Gingold H, Shaul S, and Benjamini Y. 2016. Identifying the ligated amino acid of archaeal tRNAs based on positions outside the anticodon. RNA. 22: 1477–1491.

Giegé R, Sissler M, and Florentz C. 1998. Universal rules and idiosyncratic features in tRNA identity. Nucleic Acids Res. 26: 5017.

Giegé R, Touzé E, Lorber B, Théobald-Dietrich A, and Sauter C. 2008. Crystallogenesis trends of free and liganded aminoacyl-tRNA synthetases. Crystal Growth & Design. 8: 4297–4306.

Gouy M, Guindon S, and Gascuel O. 2010. SeaView version 4: A multiplatform graphical user interface for sequence alignment and phylogenetic tree building. Molecular Biology and Evolution. 27: 221–224.

Hasegawa M, Kishino H, and Yano T. 1985. Dating of the human-ape splitting by a molecular clock of mitochondrial DNA. Journal of Molecular Evolution. 22: 160–174.

Hennig W et al. 1981. Insect Phylogeny. John Wiley & Sons Ltd.

Holbrook SR, Sussman JL, Warrant RW, Church GM, and Kim SH. 1977. RNA-ligand interactions:(I) magnesium binding sites in yeast tRNA^Phe^. Nucleic Acids Research. 4: 2811–2820.

Holley RW. 1965. Structure of an alanine transfer ribonucleic acid. Journal of the American Medical Association. 194: 868–871.

Holley RW, Apgar J, Everett GA, Madison JT, Marquisee M, Merrill SH, Penswick JR, and Zamir A. 1965. Structure of a Ribonucleic Acid. Science. 147: 1462–1465.

Huelsenbeck JP and Ronquist F. 2001. MRBAYES: Bayesian inference of phylogenetic trees. Bioinformatics. 17: 754–755.

Humphrey W, Dalke A, and Schulten K. 1996. VMD: visual molecular dynamics. Journal of Molecular Graphics. 14: 33–8–27–8.

Jack A, Ladner JE, Rhodes D, Brown RS, and Klug A. 1977. A crystallographic study of metal-binding to yeast phenylalanine transfer RNA. Journal of Molecular Biology. 111: 315–328.

Jühling F, Mörl M, Hartmann RK, Sprinzl M, Stadler PF, and Pütz J. 2009. tRNAdb 2009: compilation of tRNA sequences and tRNA genes. Nucleic Acids Research. 37: D159–62.

Kelly P, Hadi-Nezhad F, Liu DY, Lawrence TJ, Linington RG, Ibba M, and Ardell DH. 2020. Targeting tRNA-synthetase interactions towards novel therapeutic discovery against eukaryotic pathogens. PLoS Negl Trop Dis. 14: e0007983.

Kent WJ, Sugnet CW, Furey TS, Roskin KM, Pringle TH, Zahler AM, and Haussler D. 2002. The Human Genome Browser at UCSC. Genome Res. 12: 996–1006.

Lanave C, Preparata G, Saccone C, and Serio G. 1984. A new method for calculating evolutionary substitution rates. Journal of Molecular Evolution. 20: 86–93.

Laslett D and Canback B. 2004. ARAGORN, a program to detect tRNA genes and tmRNA genes in nucleotide sequences. Nucleic Acids Research. 32: 11–16.

Lawrence TJ, Amrine KC, Swingley WD, and Ardell DH. 2019. tRNA functional signatures classify plastids as late-branching cyanobacteria. BMC Evol. Biol. 19: 224.

Lowe TM and Eddy SR. 1997. tRNAscan-SE: a program for improved detection of transfer RNA genes in genomic sequence. Nucleic Acids Research. 25: 955–964.

Marck C, Grosjean H. 2002. tRNomics: analysis of tRNA genes from 50 genomes of Eukarya, Archaea, and Bacteria reveals anticodon-sparing strategies and domain-specific features. RNA 8(10):1189–232.

McQuilton P, St Pierre SE, Thurmond J, and FlyBase Consortium. 2012. FlyBase 101–the basics of navigating FlyBase. Nucleic Acids Research. 40: D706–14.

Nawrocki EP, Kolbe DL, and Eddy SR. 2009. Infernal 1.0: inference of RNA alignments. Bioinformatics. 25: 1335–1337.

Nei M and Rooney AP. 2005. Concerted and Birth-and-Death Evolution of Multigene Families. Annual Review of Genetics. 39: 121–152.

Nemenman I, Shafee F, and Bialek W 2002. Entropy and inference, revisited. English (US). In: Advances in Neural Information Processing Systems 14 - Proceedings of the 2001 Conference, NIPS 2001. Advances in Neural Information Processing Systems. Neural Information Processing Systems Foundation.

Phillips, J. B. (2013). Evolution of Eukaryotic Transfer Ribonucleic Acid. Dissertation, University of California, Merced. ProQuest ID: Phillips_ucmerced_1660D_10055. Merritt ID: ark:/13030/m5n600vj. Retrieved from https://escholarship.org/uc/item/8np368qs on January 2, 2021.

Plummer M, Best N, Cowles K, and Vines K. 2006. CODA: Convergence Diagnosis and Output Analysis for MCMC. R News. 6: 7–11.

R Core Team 2013. R: A Language and Environment for Statistical Computing. R Foundation for Statistical Computing. Vienna, Austria.

Rodríguez F, Oliver JL, Mariń A, and Medina JR. 1990. The general stochastic model of nucleotide substitution. Journal of Theoretical Biology. 142: 485–501.

Rogers HH, Bergman CM, and Griffiths-Jones S. 2010. The evolution of tRNA genes in *Drosophila*. Genome Biol Evol. 2: 467–477.

Rogers HH and Griffiths-Jones S. 2014. tRNA anticodon shifts in eukaryotic genomes. RNA. 20: 269–281.

Ronquist F, Teslenko M, Mark P van der, Ayres DL, Darling A, Höhna S, Larget B, Liu L, Suchard MA, and Huelsenbeck JP. 2012. MrBayes 3.2: efficient Bayesian phylogenetic inference and model choice across a large model space. Systematic Biology. 61: 539–542.

Sabi R, Volvovitch Daniel R, Tuller T. 2017. stAIcalc: tRNA adaptation index calculator based on species-specific weights, Bioinformatics 33: 589–591.

Saks ME, Sampson JR, and Abelson J. 1998. Evolution of a Transfer RNA Gene Through a Point Mutation in the Anticodon. Science. 279: 1665–1670.

Sarkar D. 2008. Lattice: Multivariate Data Visualization with R. Springer, New York.

Schneider TD, Stormo GD, Gold L, and Ehrenfeucht A. 1986. Information content of binding sites on nucleotide sequences. Journal of Molecular Biology. 188: 415–431.

Scott JG et al. 2014. Genome of the house fly, *Musca domestica* L., a global vector of diseases with adaptations to a septic environment. Genome Biol. 15: 466.

Sethi A, Eargle J, Black AA, and Luthey-Schulten Z. 2009. Dynamical networks in tRNA:protein complexes. Proceedings of the National Academy of Sciences of the United States of America. 106: 6620–6625.

Shi H and Moore PB. 2000. The crystal structure of yeast phenylalanine tRNA at 1.93 Å resolution: a classic structure revisited. RNA. 6: 1091–1105.

Sprinzl M, Horn C, Brown M, Ioudovitch A, and Steinberg S. 1998. Compilation of tRNA sequences and sequences of tRNA genes. Nucleic Acids Research. 26: 148–153.

Stajich JE, Block D, et al. 2002. The BioPerl toolkit: Perl modules for the life sciences. Genome Research. 12: 1611–1618.

Stajich JE and Hahn MW. 2005. Disentangling the effects of demography and selection in human history. Molecular Biology and Evolution. 22: 63–73.

Tamaki S, Tomita M, Suzuki H and Kanai A. 2018. Systematic analysis of the binding surfaces between tRNAs and their respective aminoacyl tRNA synthetase based on structural and evolutionary data. Front. Genet. 8:227.

Tamura K, Subramanian S, and Kumar S. 2003. Temporal patterns of fruit fly (*Drosophila*) evolution revealed by mutation clocks. Molecular Biology and Evolution. 21: 36–44.

Tåquist H, Cui Y, and Ardell DH. 2007. TFAM 1.0: an online tRNA function classifier. Nucleic Acids Research. 35: W350–3.

Tavaré S. 1986. Some probabilistic and statistical problems in the analysis of DNA sequences. Lectures in Mathematics of Life Sciences. 17: 57–86.

Tremblay-Savard O, Benzaid B, Lang BF, and El-Mabrouk N. 2015. Evolution of tRNA repertoires in *Bacillus* inferred with OrthoAlign. Mol Biol Evol. 32:

Tweedie S et al. 2009. FlyBase: enhancing *Drosophila* Gene Ontology annotations. Nucleic Acids Res. 37: D555–D559.

Velandia-Huerto CA, Berkemer SJ, Hoffmann A, Retzlaff N, Romero Marroquín LC, Hernández-Rosales M, Stadler PF, and Bermúdez-Santana CI. 2016. Orthologs, turn-over, and remolding of tRNAs in primates and fruit flies. BMC Genomics. 17: 617.

Wiegmann BM, Yeates DK, Thorne JL, and Kishino H. 2003. Time flies, a new molecular time-scale for brachyceran fly evolution without a clock. Systematic Biology. 52: 745–756.

Wolfson AD, LaRiviere FJ, Pleiss JA, Dale T, Asahara H, and Uhlenbeck OC 2001. tRNA Conformity. In: The Ribosome. CSHL Press, pp. 185–193.

Zamudio GS, José MV. 2018. Identity elements of tRNA as derived from information analysis. Orig Life Evol Biosph. 48(1):73–81.

