## Supplementary material for "Structural and Genetic Determinants of Convergence in the *Drosophila* tRNA Structure-Function Map": SOM for Phillips and Ardell 2021

### Supplementary Online Materials for Phillips and Ardell "Structural and Genetic Determinants of Convergence in the *Drosophila* tRNA Structure-Function Map"

#### CONTENTS


1 Code and Data 1

2 Substitutions in Ion-Binding Pocket by Functional Class and Location 1

3 Substitutions in Ion-Binding Pockets by Chromosomal Location 3

4 Transition and Transversion Rates in *Drosophila* tRNA Genes 3

5 Site Divergences with Other Evolutionary Models 3

6 Significance of Paired-Site CIFs in *D. melanogaster* 3

7 Average Frequency and Information of Nucleotides in tRNA genes in *D. melanogaster* 3


#### 1. CODE AND DATA

The full code and data to reproduce all results is available at http://dx.doi.org/10.6084/m9.figshare.12713705.

#### 2. SUBSTITUTIONS IN ION-BINDING POCKET BY FUNCTIONAL CLASS AND LOCATION

**Table S1.** Sequence patterns, ortholog sets, functional classes and locations (in *D. melanogaster*) contributing to elevated substitution rates in an ion-binding pocket. Results are provided for the largest subset of data partitions (X indicates the species is excluded from that partition). Species bit-string corresponds in order to: *D. melanogaster, D. simulans, D. sechellia, D. yakuba, D. erecta, D.ananassae, D pseudoobscura, D. persimilis, D. willistoni, D. mojavensis, D. virilis, D. grimshawi*. Functional classes use IUPAC amino acid one-letter abbreviations for elongator tRNAs, "X" for initiator tRNAs, "U" for selenocysteine tRNAs.

| Species Bit-String | Coordinate(s) | Ortholog Set (Class Anticodon):Sequence | Location in ***D. melanogaster*** |
|---|---|---|---|
| 111111110111 | 16:17 | 72 (U TCA):TGGGGGGG0GGG:<br>-TTTTTTTTTT | 2R:c(7245069..7245155) |
| 111111110111 | 16:60 | 2 (R TCT):TTTTTCCC0TCC:<br>TTTTTCCCTCC | 2L:1965426..1965498 |
| | | 92 (L AAG):CCCCCCCC0CCT<br>112(I AAT):TTTTTTTT0CCC<br>114(X CAT):TTTTTTCT0TTT | 2R:c(10871832..10871913)<br>2R:15603070..15603143<br>2R:15613410..15613481 |
| 111111110111 | 16 | | |

**Table S1 – continued from previous page**

| Species Bit-String | Coordinate(s) | Ortholog Set (Class Anticodon):Sequence | Location in *D. melanogaster* |
|---|---|---|---|
| | | 136(S TGA):TTTTTCTT0TTT | 2R:c(18959542..18959623) |
| | | 137(S TGA):TTTTTCTT0TTT | 2R:18960104..18960185 |
| | | 188(P CGG):AAAAAAAA0GGG | 3L:18611490..18611561 |
| | | 192(R TCG):AAAAAAAA0TTT | 3R:1213950..1214022 |
| | | 226(V AAC):TTTTTTCT0TTT | 3R:12147412..12147484 |
| | | 251(V CAC):CCCCCCTC0TCC | 3R:15615844..15615916 |
| | | 285(S AGA):CCCCCCCC0ACC | X:c(13919142..13919223) |
| 111111110111 | 17 | 46 (N GTT):TTTTTCTT0CCT | 2R:c(2040108..2040181) |
| | | 77 (M CAT):TTTTTTTT0ATT | 2R:c(7548068..7548140) |
| 111111110111 | 17 | 102(F GAA):T-TTTTTT0TTT | 2R:c(13492040..13492112) |
| | | 138(N GTT):TTTTTCTT0CCT | 2R:20261586..20261659 |
| | | 186(M CAT):TTTTTTTT0AAA | 3L:16343017..16343089 |
| | | 205(N GTT):TTTTTCTT0CCT | 3R:c(3965255..3965328) |
| 111111110111 | 60 | 231(A AGC):GGGGGAGG0GGG | 3R:13445460..13445532 |
| | | 240(A AGC):GGGGGAGG0GGG | 3R:c(13471029..13471101) |
| | | 245(A AGC):GGGAGAGG0AGG | 3R:c(13484103..13484175) |
| | | 246(A AGC):GGGGGAGG0AGG | 3R:13493909..13493981 |
| | | 252(A AGC):GGGGGAGG0GGG | 3R:15616694..15616766 |
| 11111111011X | 16 | 260(S GCT):TTTTTATT0AAX | 3R:c(18222902..18222983) |
| 11111111011X | 60 | 221(T TGT):CCCCCCTT0TCX | 3R:c(8148285..8148356) |
| | | 242(A AGC):GGGGGAGG0GGX | 3R:13472090..13472162 |
| 1X111111011X | 16 | 258(S GCT):AXAAATAA0TTX | 3R:18222250..18222331 |
| | | 259(S GCT):AXAAAAAA0TTX | 3R:c(18222521..18222602) |
| 111111110XXX | 16 | 83 (I AAT):TCCCTTCC0XXX | 2R:c(9317787..9317860) |
| | | 85 (I AAT):TTTCTTCC0XXX | 2R:9318489..9318562 |
| | | 272(Q TTG):CCCTTTTT0XXX | X:c(3321485..3321556) |
| 111111110XXX | 17 | 193(M CAT):TTTTTAAA0XXX | 3R:c(2321761..2321833) |
| 111111110XXX | 60 | 234(A AGC):GGGGGAGG0XXX | 3R:13448268..13448340 |
| | | 238(A AGC):GGGGGAGG0XXX | 3R:c(13456764..13456836) |
| | | 244(A AGC):AGGGGAGG0XXX | 3R:13482867..13482939 |
| 11XX111X0111 | 16 | 287(S AGA):TTXXTTCX0CCC | X:13964674..13964755 |
| 11X1111X01X1 | 16 | 292(P CGG):AAXAAGTX0TXT | X:18459797..18459868 |
| 1111X1110XXX | 17 | 61 (N GTT):TTTTXCTT0XXX | 2R:c(2077634..2077707) |
| 1XXX111X0111 | 60 | 166(A AGC):GXXXGAGX0GGG | 3L:c(8021646..8021718) |
| 111111X10XXX | 16 | 273(Q CTG):TTTTTTXC0XXX | X:3713732..3713803 |
| | | 294(I AAT):TTTTTTXC0XXX | X:c(21102352..21102426) |
| 1XX1111X01X1 | 16 | 220(T TGT):TXXTTTTX0TXA | 3R:8032297..8032368 |

**Table S1 – continued from previous page**

| Species Bit-String | Coordinate(s) | Ortholog Set (Class Anticodon):Sequence | Location in *D. melanogaster* |
|---|---|---|---|
| 1X1111110XXX | 16 | 274(P CGG):GXGAAGTT0XXX | X:3721655..3721726 |
| | | 289(R TCG):TXTTTTCC0XXX | X:c(13997752..13997824) |
| 1X1111110XXX | 17 | 47 (N GTT):TXTTTCTT0XXX | 2R:2040691..2040764 |

#### 3. SUBSTITUTIONS IN ION-BINDING POCKETS BY CHROMOSOMAL LOCATION

#### 4. TRANSITION AND TRANSVERSION RATES IN *DROSOPHILA* TRNA GENES

#### 5. SITE DIVERGENCES WITH OTHER EVOLUTIONARY MODELS

We ran analogous site divergence calculations using the HKY model (Hasegawa, M., H. Kishino, and T. Yano (1985). Dating of the human-ape splitting by a molecular clock of mitochondrial DNA. *Journal of Molecular Evolution* 22(2), 160–174) and the HKY+Γ model. We estimated substitution rates with MrBayes 3.2.1 (Huelsenbeck and Ronquist 2001; Ronquist et al. 2012) using the fixed known species tree (Drosophila 12 Genomes Consortium 2007). For all runs we constrained change to the tree topology by setting rates of stochastic TBR and branch multipliers to zero probability. All Bayesian analyses were run with two simultaneous chains for $4 \times 10^6$ iterations, monitoring convergence of split frequency standard deviations and saved parameters every 500 iterations.

#### 6. SIGNIFICANCE OF PAIRED-SITE CIFS IN *D. MELANOGASTER*

#### 7. AVERAGE FREQUENCY AND INFORMATION OF NUCLEOTIDES IN TRNA GENES IN *D. MELANOGASTER*

**Table S2.** Sequence changes in ion-binding pocket sites by chromosome (Muller elements A – E), when location data was available for *D. melanogaster*. The X chromosome (Muller element A) has a larger fraction of change compared to other chromosome arms.

| Chromosome Location | Ortholog Sets with Changes in Sites 15, 18, 19, 20, or 59 (% of total) | Ortholog Sets with Changes in Sites 16, 17 or 60 (% of total) |
|---|---|---|
| 2L | 5 (0.119) | 4 (0.095) |
| 2R | 10 (0.096) | 23 (0.221) |
| 3L | 6 (0.113) | 7 (0.132) |
| 3R | 6 (0.073) | 21 (0.256) |
| X | 9 (0.273) | 11 (0.333) |
| Not Available | 14 (0.030) | 18 (0.039) |

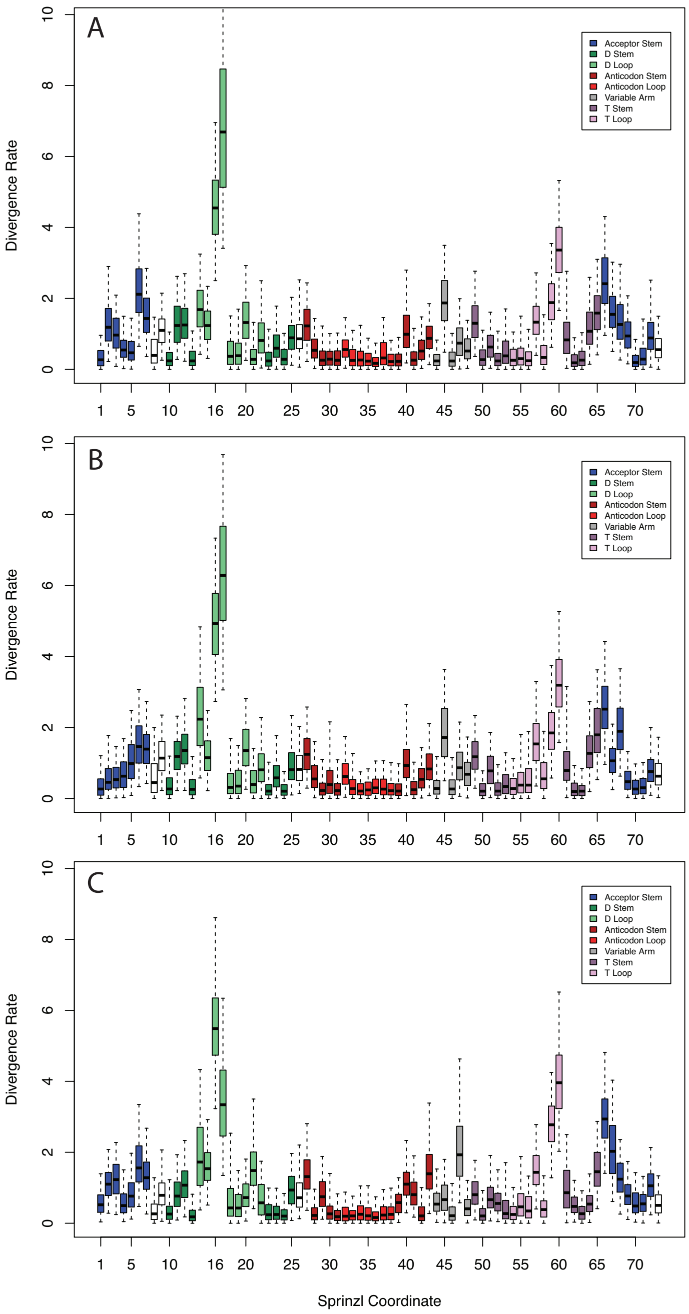


**Fig. S1.** Site divergence rates with a HKY evolutionary model. (A) the set with the largest number of species (111111110111, missing *D. willistoni*); 84 orthologous sets); (B) the set with the second largest tree length (110011100111, second only to the set with largest number of species, set contains *D. melanogaster, D. simulans, D. erecta, D.ananassae, D pseudoobscura, D. mojavensis, D. virilis, D. grimshawi*); 86 orthologous sets; and (C) the set with the largest number of included ortholog sets (101111110000, *D. melanogaster, D. sechellia, D. yakuba, D. erecta, D.ananassae, D. pseudoobscura, D. persimilis*); 155 ortholog sets

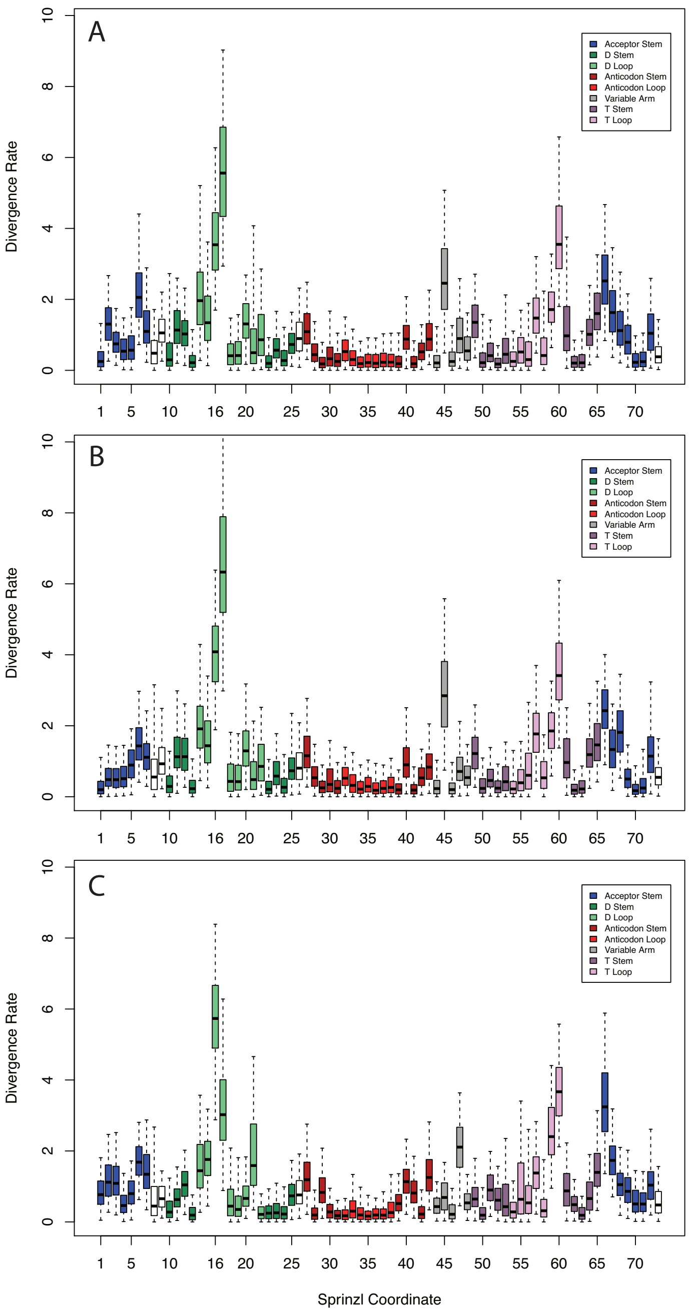


**Fig. S2.** Site divergence rates for three Mr.Bayes analyses show confirmatory results to the GTR model when simulations are run with a HKY+$\gamma$ evolutionary model. Sites show similar patterns to the structural divergence rates in the GTR model data.. (A) the set with the largest number of species (111111110111, missing *D. willistoni*); 84 orthologous sets); (B) the set with the second largest tree length (110011100111, second only to the set with largest number of species, set contains *D. melanogaster, D. simulans, D. erecta, D.ananassae, D pseudoobscura, D. mojavensis, D. virilis, D. grimshawi*); 86 orthologous sets; and (C) the set with the largest number of included ortholog sets (101111110000, *D. melanogaster, D. sechellia, D. yakuba, D. erecta, D.ananassae, D. pseudoobscura, D. persimilis*); 155 ortholog sets

**Table S3.** Median and Credible Intervals of Transition and Transversion Rates in *Drosophila* tRNA Genes

| | **101111110000** | **110011100111** | **111111110111** |
|---|---|---|---|
| **C↔T** | 0.2726 (0.1725, 0.3756) | 0.2491 (0.1402, 0.3657) | 0.2662 (0.1979, 0.3875) |
| **A↔G** | 0.4828 (0.3557, 0.6201) | 0.4947 (0.3325, 0.6455) | 0.4889 (0.4031, 0.6464) |
| **A↔C** | 0.0456 (0.0053, 0.0942) | 0.1045 (0.0353, 0.1951) | 0.1047 (0.0510, 0.1857) |
| **G↔T** | 0.0676 (0.0250, 0.1140) | 0.0508 (0.0147, 0.0914) | 0.0472 (0.0224, 0.0873) |
| **A↔T** | 0.0600 (0.0191, 0.1078) | 0.0427 (0.0048, 0.0882) | 0.0426 (0.0158, 0.0878) |
| **C↔G** | 0.0714 (0.0297, 0.1161) | 0.0583 (0.0192, 0.1036) | 0.0504 (0.0286, 0.0912) |

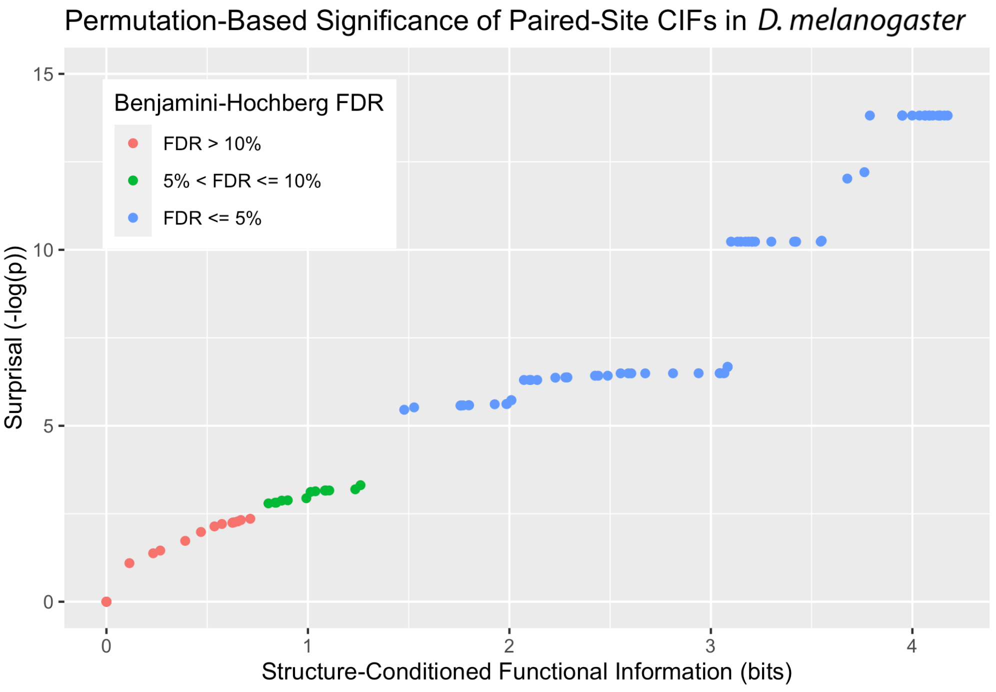


**Fig. S3.** The Surprisal of paired-site tRNA CIFs in *D. melanogaster* as a function of the total structure-conditioned functional information (stack-heights) of CIFs, and colored by computed Benjamini-Hochberg FDRs, as computed by tSFM v1.0.

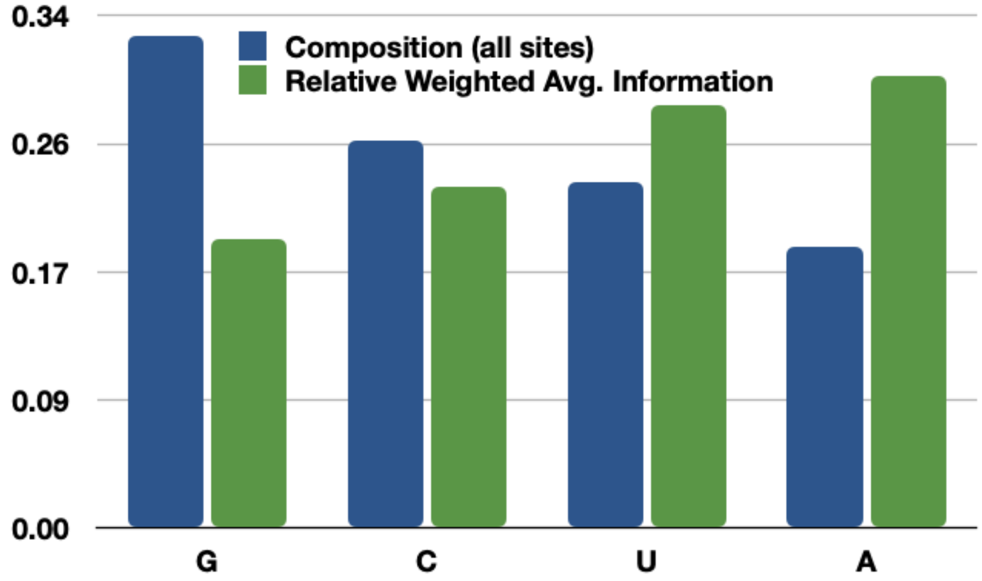


**Fig. S4.** Average composition and relative average structure-conditioned functional information of the four RNA nucleotides in 288 *Drosophila* tRNA genes with COVE bit-scores $\geq$ 50 bits.
